# Spike transmission failures in axons from mouse cortical pyramidal neurons in vivo

**DOI:** 10.1101/2024.01.29.577733

**Authors:** Netanel Ofer, Victor Hugo Cornejo, Rafael Yuste

**Author notes:** **For correspondence:** (RY). Edmond and Lily Safra Center for Brain Sciences, The Hebrew University of Jerusalem, Jerusalem, Israel (Netanel Ofer). Facultad de Ciencias Biológicas, Universidad Católica, Santiago, Chile (Victor Hugo Cornejo).

## Abstract

The propagation of action potentials along axons is traditionally considered to be reliable, as a consequence of the high safety factor of action potential propagation. However, numerical simulations have suggested that, at high frequencies, spikes could fail to invade distal axonal branches. Given the complex morphologies of axonal trees, with extensive branching and long-distance projections, spike propagation failures could be functionally important. To explore this experimentally *in vivo*, we used an axonal-targeted calcium indicator to image action potentials at axonal terminal branches in superficial layers from mouse somatosensory cortical pyramidal neurons. We activated axons with an extracellular electrode, varying stimulation frequencies, and computationally extracted axonal morphologies and associated calcium responses. We find that axonal boutons have higher calcium accumulations than their parent axons, as was reported *in vitro*. But, contrary to previous *in vitro* results, our data reveal spike failures in a significant subset of branches, as a function of branching geometry and spike frequency. The filtering is correlated with the geometric ratio of the branch diameters, as expected by cable theory. These findings suggest that axonal morphologies contribute to signal processing in the cortex.

## Introduction

In the cerebral cortex, axons of pyramidal cells can be very complex with long-distance projections, including intrinsic collaterals with arbors in different cortical layers, extrinsic projections across cortical areas and between brain regions, and callosal connections (***Rockland, 2020***). Whether all action potentials propagate faithfully throughout these anatomically complex axonal arbors has long been debated (***Raastad and Shepherd, 2003***). Experiments *in vitro* have shown reliable transmission of individual spikes and spike trains through the axonal arbor of cortical pyramidal neurons (***Cox et al., 2000***; ***Koester and Sakmann, 2000***; ***Popovic et al., 2011***). However, cable theory analyses predict that geometrical heterogeneities, such as changes in axonal diameter or bifurcation points, may cause a delay or even a failure of an action potential propagation as a result of the different electrical impedance at these points (***Goldstein and Rall, 1974***; ***Manor et al., 1991***). Indeed, numerical simulations of action potential trains at branching points have shown differential propagation of firing patterns as a function of axonal branch diameters and spike frequency, suggesting differential neuronal information processing by specific axonal branches (***Ofer et al., 2017, 2020***). Thus, there is a discrepancy between experimental results and theoretical predictions. One possibility is that the reported fidelity of action potential propagation at branching points and in distal axons could be ensured by non-uniform densities of sodium and potassium voltage-gated channels, which compensate for morphological changes (***Cho et al., 2017***; ***Sabater et al., 2021***; ***Zang and Marder, 2021***). Moreover, since these experimental data was taken *in vitro*, it is possible the axonal propagation could be different *in vivo*, since many physiological factors could influence propagation and alter the outcome.

To measure spike propagation *in vivo* we used two-photon imaging of action potentials with calcium indicators (***Yuste and Denk, 1995***). In contrast to electrophysiology, this optical approach allows the measurement of the signal throughout neuronal processes – including fine dendritic and axonal branches. Despite the relatively slow temporal resolution of calcium imaging, it can detect action potential activity in pyramidal cells at the single-spike level (***Smetters et al., 1999***; ***Chen et al., 2013***). Advanced genetically encoded voltage indicators (GEVI), are also capable of reporting axonal activity with single action potential resolution (***Cornejo et al., 2022***), but their application is still challenging due to the low signal to noise and high sampling rate required in imaging large regions of interest (ROIs) along axonal branches. To specifically record action potential activity in axons, one can use axon-targeted GCaMP6s, characterized by a uniform expression and distribution, sufficient brightness, high signal-to-noise ratio, and photostability (***Broussard et al., 2018***). To capture responses to dozens of spikes at high frequency, GCaMP6s actually outperforms the newer jGCaMP8, owing to its larger dynamic range with more linearity and less saturation (***Zhang and Looger, 2023***). Because of this, to monitor activity across axonal branching points in response to high-frequency action potential trains *in vivo*, we imaged axon-targeted GCaMP6s expressed in superficial axonal branches of pyramidal cells from mouse primary somatosensory cortex *in vivo*, in response to extracellular electrical stimulation. We aimed to determine whether spikes originating from an axonal parent branch successfully reached the daughter branches. We found a heterogeneity of responses. In most axons (11 out of 17), spikes propagated from the parent branch into both daughter branches reliably, resulting in a similar response at all branches. But in 6 cases, higher frequency spikes failed at the bifurcation point, leading to different responses in the daughter branches. As predicted by cable theory, morphological analysis of these cases revealed a correlation between the geometrical ratio (GR) of the parent and daughter diameters and the reliability of spike transmission, which also decreased with the frequency of the action potential trains. Our results reveal that action potentials fail to propagate in a significant number of axonal branches in mouse cortical pyramidal neurons *in vivo*.

## Methods

### Animal use and care

All experimental procedures in animals were approved by the Columbia University Institutional Animal Care and Use Committee (IACUC, protocol AC-AABN3562) in compliance with the National Institutes of Health guidelines for the care and use of laboratory animals. We used both female and male wild-type C57BL/6 that were 2-4 months of age, kept on a continuous 12-hour light/dark cycle, and freely available for food and water.

### Viral injections surgeries

Mice were anesthetized with 2% isoflurane and placed in a stereotaxic atop a heating pad maintained at 37°C. Enrofloxacin (5 mg/kg) and carprofen (5 mg/kg) were injected intraperitoneal and lidocaine (2 mg/kg) subcutaneously in the site of the skin incision over the midline of the scalp. A 0.5 mm hole was drilled in the skull over the anterior left portion of the somatosensory cortex. Virus pAAV-hSynapsin1-axon-GCaMP6s-P2A-mRuby3 (Addgene viral prep #1120050-AAV5) was injected into layer 2/3 of the primary somatosensory barrel (S1BF) cortex (3.2 mm lateral, 0.1 mm posterior and 1.6 mm down from the bregma) to target intra-cortical axons of pyramidal cells. 150 nl of the viral prep was injected at a rate of 50 nl/min, and 3 min wait period before needle withdrawal.

### Head plate and cranial windows implantation

After 2-3 weeks of viral injection surgeries, mice were anesthetized with 2% isoflurane in the same stereotaxic rig, and a titanium head plate was attached to the skull using dental cement. A 3 mm round craniotomy was made over the primary somatosensory cortex, and the dura was removed. The upper tangent of the 3 mm round glass coverslip (Warner Instruments, CS-3R) was placed at the same site as the viral injection and fixed to the skull using cyanoacrylate.

### Extracellular electrode stimulation and two-photon imaging

Immediately after head plate and cranial window implantation, the animal was moved to a twophoton microscope rig, and a concentric bipolar electrode of 200/50 *μ*m diameter (outer/inner pole) (FHC, #30215) was placed at the same coordinates as the viral injection site in the somatosensory cortex. Stimulation was performed with a stimulus isolator (World Precision Instruments, A365) in bipolar mode. 1 ms pulses were generated using a Master-8 pulse generator (A.M.P.I.) to the isolator. Axonal branch searching was performed by giving a test pulse consisting of 5 pulses at 50 Hz, with the isolator set to 100 *μ*A. Once target axonal branches were found, minimal stimulation was tested to observe significant responses, and the isolator was set in a range of 40 to 100 *μ*A.

*In vivo* imaging was performed in layers 1 and 2/3 at 50–150 *μ*m below the cortical surface of the exposed mouse cortex with a custom two-photon microscope (adapted from Prairie model, Bruker), with a 25x/1.05 N.A. water immersion objective (Olympus) and a tunable Ti-sapphire laser (Mai Tai eHP DS, Spectra-Physics). Animals expressing axon-GCAMP6s in the cortex were imaged with the two-photon system with the laser tuned at 940 nm and signals were collected with filters 510/20 nm for GCAMP6s and 605/15 for mRuby3. Axons were imaged at a resolution of 256x256 pixels to 10 or 12x zoom at 60 Hz (pixel size of 0.166 *μ*m or 0.2 *μ*m, accordingly). Imaging power laser at 940 nm was measured at the end of the objective and for regular imaging, 50-80 mW was used for all experiments.

### Electrical stimulation

Six different series of electrical pulses with frequencies of 40, 60, 80, 100, 120, and 140 Hz were injected for a constant duration of 200 ms, which resulted in sequences of 8, 12, 16, 20, 24, and 28 pulses. The 1 ms pulses were designed in a way that each pulse led to a single action potential. The time interval between pulse series was 7 or 8 seconds to allow complete decay of the fluorescence before the next series of pulses. The stimulus for each frequency was repeated seven times in random order (Figure 1A). The stimulus was designed in that way, with a different number of spikes for each frequency, because of the nonlinearity relationship between calcium concentration and the observed fluorescence. This nonlinearity is due to the properties of the indicator, which requires binding of four calcium ions for lighting, causing supralinear and sublinear regimes that can be approximated by the Hill equation (***Pologruto et al., 2004***). To address the challenge of inferring the number of spikes from the fluorescence signal, we employed a strategy wherein the fluorescence signal from one branch at lower frequencies was used as a baseline to estimate the number of spikes that failed to propagate in another branch at higher frequencies.

**Figure 1.**
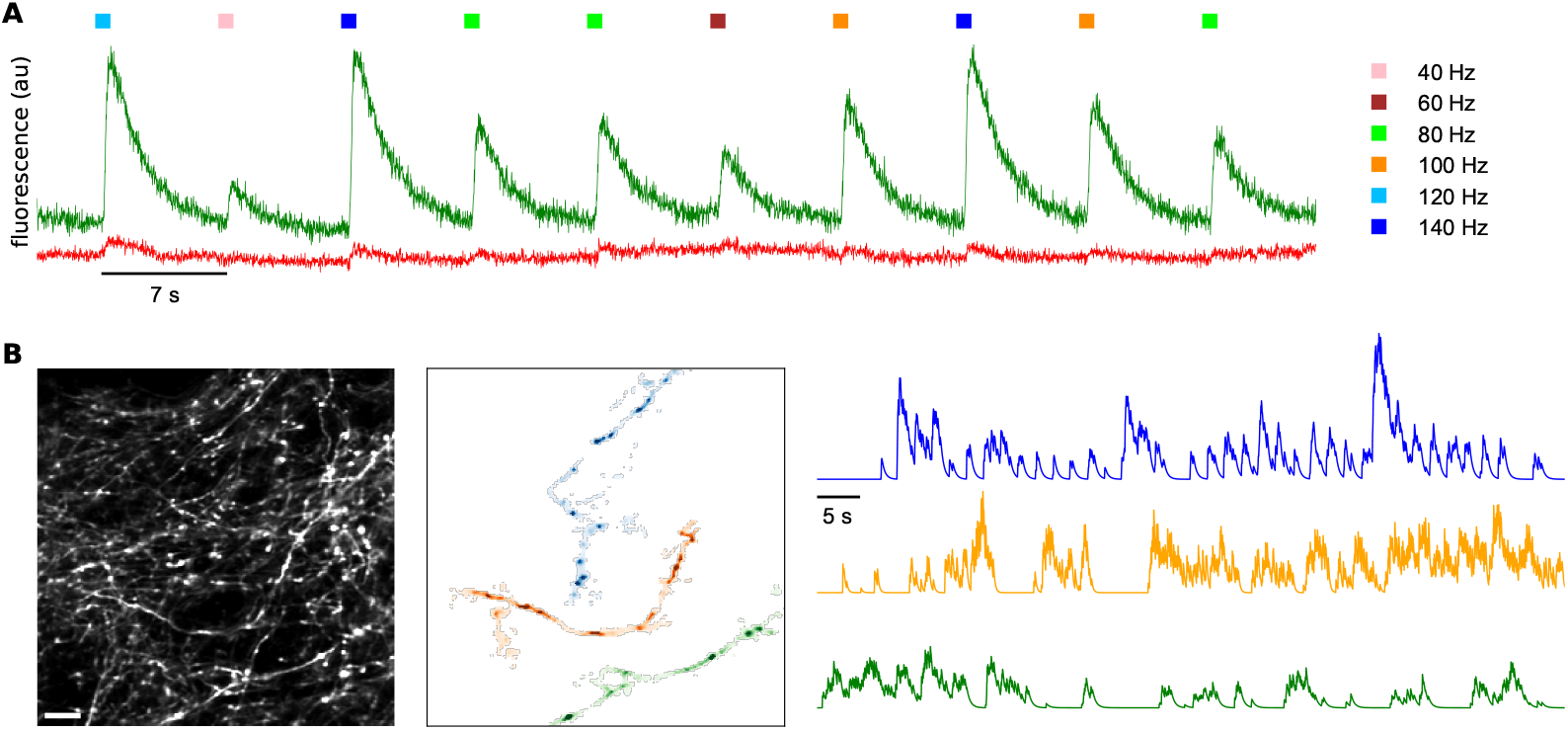
Experimental design and spatiotemporal analysis of axonal branches. A: Fluorescence responses of axons to electrical stimulation of neuropil at different stimulation frequencies; GCaMP6s in green, mRuby3 in red. B: Computational extraction of axonal branches, separating different axons and neuropil background based on activity. Scale bar: 10 *μ*m.

### Image analysis

Motion correction was applied on the GCaMP6s channel using *CaImAn* (***Giovannucci et al., 2019***). The same correction in X and Y that was conducted on the GCaMP6s channel was applied to the mRuby3 channel. Then, the GCaMP6s channel was normalized by the mRuby3 channel to correct axonal branches at a different distance from the focal plane. We used *CaImAn* to make masks only for the axonal branched of the bifurcation points (Figure 1B). The *‘SparseNMF’* initialization strategy was used for quickly uncovering spatial structure in the imaging data, especially for neural processes such as dendrites or axons, where the degree of overlap between the different branches is higher (***Pnevmatikakis et al., 2016***). Then, we dissected the ROIs of the parent and the two daughter branches (see Figure 2B). To know which of the branches is the parent branch, which the spikes are coming from, we compared the time of the response at three small ROIs at the farthest edges of the branches. The branch with the earliest onset of the signal is considered to be the parent branch. Sometimes it is evident which branch is the parent branch, for example, where the bifurcation is in a ‘Y’ shape. However, in some cases, we cannot be sure which branch is the parent branch based only on morphology.

**Figure 2.**
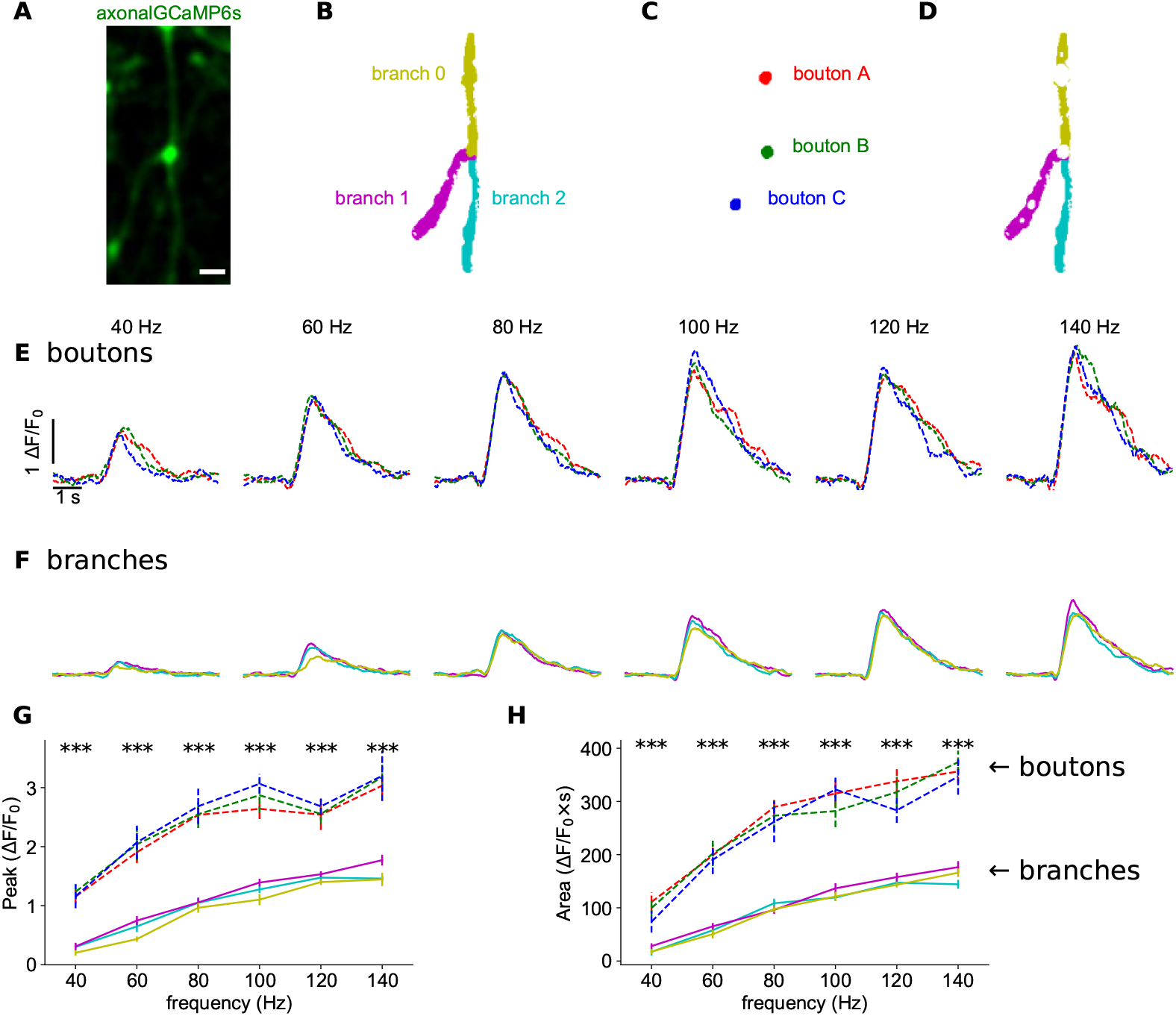
Increased calcium responses at axonal boutons. A: Time-averaged image of GCaMP6s activity during electrical stimulation. Scale bar: 2 *μ*m. B: Color map of parent (0) and two secondary branches (1 and 2). C: Masks of three axonal boutons (a, b, and c). D: Masks of axonal branches after removing the boutons. E: Normalized GCaMP6s/mRuby3 signal for each bouton at different frequency. Average of 7 trials. Colors according to boutons in 2C. F: Normalized GCaMP6s/mRuby3 signal for each branch at different frequency. Average of 7 trials. Colors according to branches in 2D. G: Peak amplitudes of signals in 2E and 2F as a function of frequency. H: Area under curve of signals in 2E and 2F as a function of frequency. Vertical lines show standard error. Asterisks indicate statistical significance difference between signals; Kruskal-Wallis H-test.

The fluorescence signals for the 7 trials were averaged separately for each branch and then were filtered with a Savitzky-Golay filter (filter length of 51; order of the polynomial was 3).

The Δ*F/F*_0_ was calculated according to Equation 1. *F*_0_ was calculated by averaging 100 frames (∼ 1.6 s) before the onset of each pulse series.

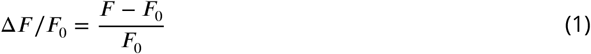

### Measurement of axonal diameters

To measure the diameters of the axonal branches, we used *ImageJ* to draw a line (width of 10 pixels) perpendicular to the axon. The fluorescence intensity pattern was fitted to the Lorentzian function, and then the full width at half maximum (FWHM) was calculated. When possible, the axonal diameter measurements were done from the mRuby3 channel and not from the GCaMP6s channel, which may be affected by the axonal activity.

### Statistical analysis

A factorial non-parametric Kruskal-Wallis H-test or two-sided unpaired Student’s t-test was applied. The correlation between parameters was examined by the Wald test with t-distribution of the test statistic. The two-sided p-value for a hypothesis test whose null hypothesis is that the slope is zero. The asterisks indicate statistical significance: n.s, not significant, *p < 0.05, **p < 0.01, ***p < 0.001.

### Data availability

Videos of the activity at the branching points are publicly available at the Columbia University Academic Commons site (https://doi.org/doi:10.7916/y615-0e51). Jupyter notebooks of the Python codes are publicly available on the Columbia University Neurotechnology Center’s GitHub page (https://github.com/NTCColumbia/).

## Results

We measured axonal calcium dynamics in 17 axonal branchpoints, likely belonging to 17 different neocortical pyramidal neurons of mouse primary somatosensory cortex, imaged in 5 different mice (Table 1). To study the propagation of action potential trains through these axonal branch-points, we injected extracellular current pulses at various frequencies into the neighboring neuropil and examined axonal responses at branch bifurcations. Stimuli were designed to produce action potential trains at six different frequencies, between 40 and 140 Hz, with a duration of 200 ms. The fluorescence of axons that expressed axon-GCaMP6s was imaged with a two-photon microscope. We searched for axons where branches in the same focal plane responded to test pulses from the electrode and imaged their responses. To correct for differences in expression and focal plane, GCaMP6s fluorescence was normalized by mRuby3 fluorescence, which was bicistronically co-expressed (Figure 1A). For each frequency, fluorescence signals of 7 trials were averaged, independently for each axonal branch – a “parent” and two “secondary” branches. The classification of branches into parent or secondary was due to their morphologies, whereby larger parent branches split into two smaller branches at an acute angle. We analyzed both the peak of the calcium signal and the area under the fluorescence curve, which respectively represent the peak current and the calcium ions charge injected. Peak signals were more sensitive to noise compared to areas, due to sampling rate and the filters applied during the analysis.

**Table 1.** Neurons used in the study.

|  | Mouse 1 | Mouse 2 | Mouse 3 | Mouse 4 | Mouse 5 | Total |
| --- | --- | --- | --- | --- | --- | --- |
| Neurons | 11 | 2 | 2 | 1 | 1 | 17 |
| Propagation | 9 | 0 | 0 | 1 | 1 | 11 |
| Failures | 2 | 2 | 2 | 0 | 0 | 6 |

### Axonal boutons generate increased calcium responses ***in vivo***

In our data, we noticed axonal boutons, both *en passant* and terminal boutons (Figure 2A). In past studies *in vitro*, it has been reported that boutons have higher peak calcium accumulations than the axonal branches themselves (***Cox et al., 2000***; ***Koester and Sakmann, 2000***). To analyze this, in a pilot experiment, we digitally extracted these boutons and conducted separate analyses of the signal at the axonal boutons (Figure 2C) and axonal branches (Figure 2D). In this experiment, the intensity of the signal in axonal boutons was higher than in the axonal branches in both peak amplitudes (2.33±0.71; mean ± std, maximum p<0.0004; Wallis H-test, Figure 2G) and area under the curves (2.75±0.67; mean ± std, maximum p<0.0003; Wallis H-test, Figure 2H). Similar results were found in other experiments, bouton/axon ratio of signal amplitudes was 3.46±1.23 (mean ± std), and the area under the curve had a ratio of 3.29±1.07 (mean ± std, n=19 boutons, 7 neurons from 7 mice). In 6 out of 7 experiments, the differences in peaks and areas between boutons and branches were statistically significant across all stimulus frequencies (maximum p<0.04; Wallis H-test), while in the last experiment, significance was observed only at higher frequencies of 100, 120, and 140 Hz (maximum p<0.0003; Wallis H-test). The higher amplitude of the responses at axonal boutons (maximum of 3.18±0.21 Δ*F/F*_0_*s* compared to 1.77±0.08 Δ*F/F*_0_*s* at the axonal branch; Figure 2G) also indicated that the fluorescence signal at the axonal branches was below the saturation regime. We concluded that axonal boutons had increased calcium accumulations than axonal branches, confirming the *in vitro* results (***Cox et al., 2000***; ***Koester and Sakmann, 2000***).

### Reliable propagation of action potentials in most axonal branches

Due to variability between boutons, and to avoid fluorescence contamination by the stronger signals from axonal boutons, in the rest of the study, we only analyzed signal from axonal branches, computationally removing the boutons during the analysis process. In most axonal bifurcations analyzed, we found similar responses in all axonal branches (Figure 3; Table 1). As the stimulation frequency increased (and presumably the number of axonal spikes), the fluorescence response became stronger (Figures 3C, 3D), but, at each frequency, the peak amplitude and area under the curve were similar in all parent and secondary branches (Figure 3D). There was no statistically significant difference between branches at any frequency, in both peak amplitude of the signal (7 trials, 6 frequencies, minimum p>0.18; Wallis H-test; Figure 3E) or area under the curve (7 trials, 6 frequencies, minimum p>0.26; Wallis H-test; Figure 3F). Similar results were obtained in 10 other cases (see below; Table 1; Figure 3 - Figure Supplement 1). These results are consistent with the possibility that the same number of action potentials in the parent branch propagated into the two secondary branches without transmission failures. We concluded that, for the majority of cases examined, axonal propagation was reliable (Figure 3G).

**Figure 3.**
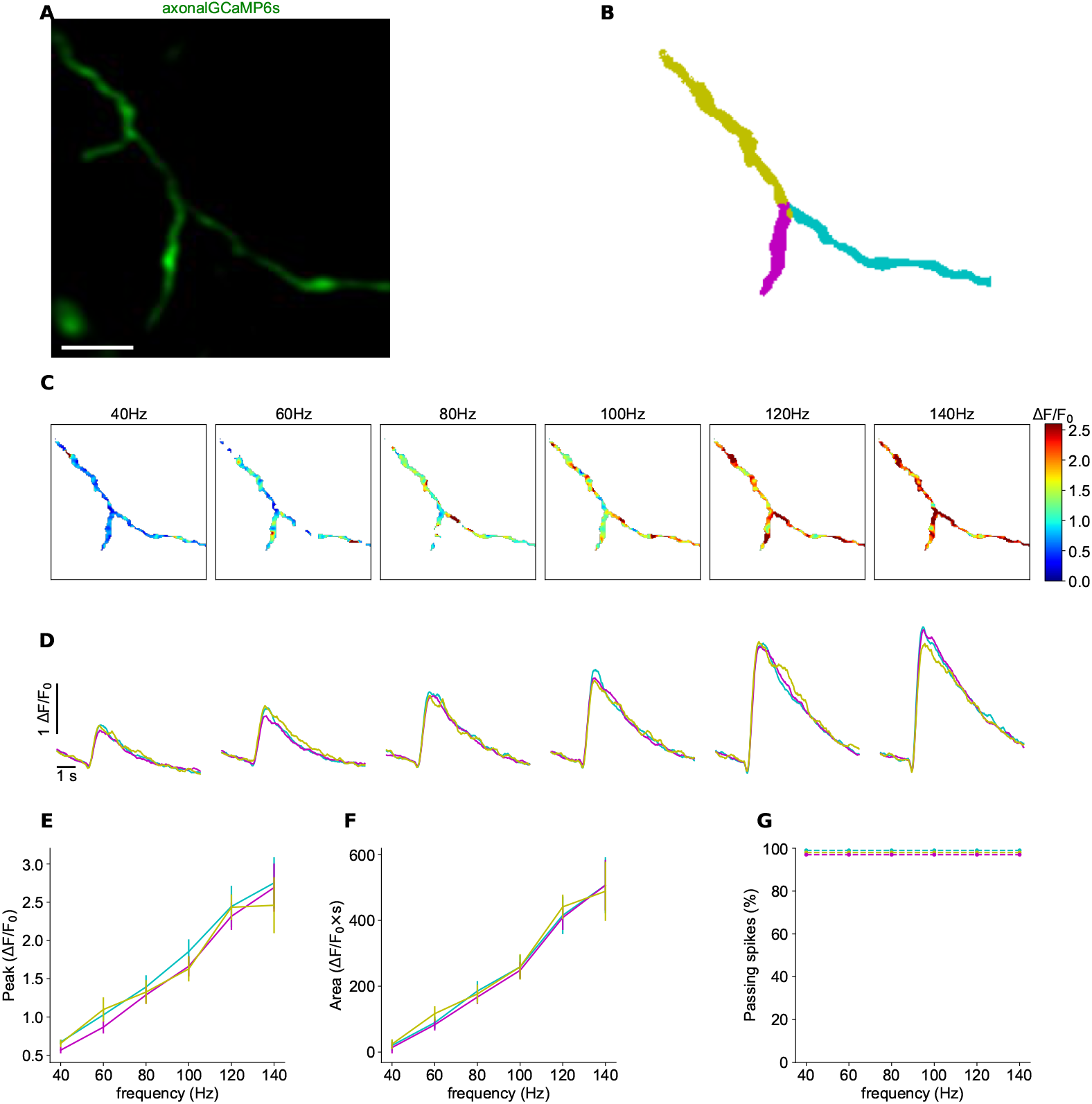
Reliable propagation of action potentials at axonal branching point. A: Time-averaged image of GCaMP6s activity during electrical stimulation. Scale bar: 5 *μ*m. B: Color map of parent (0) and two secondary branches (1 and 2). C: Maps of normalized peak amplitudes at each frequency. D: Normalized GCaMP6s/mRuby3 signal for each branch at different frequency. Average of 5 trials. Colors according to branches in 1B. E: Peak amplitudes of signals in 3D as a function of frequency. F: Area under curve of signals in 3D as a function of frequency. Vertical lines show standard error. G: Percentage of signals that propagate at each branch, as a function of frequency. Figure 3—figure supplement 1. Spike propagation at different branchpoints.

### Different responses at axonal branches

In some experiments, we also found spike filtering at a branching point where calcium signals were different between parent and secondary branches (Figure 4; Table 1). The GCaMP6s signal (Figure 4A) was normalized by the mRuby3 signal. Subsequently, branch masks were created (Figure 4B), and axonal boutons were removed (Figure 4C). At 40, 60, and 80 Hz, the fluorescence intensity was similar in all branches (n=7, significant p-value at a non-consecutive frequency in the peak signals; minimum p>0.08 for the area under the signals; Wallis H-test). However, at 100, 120, and 140 Hz, some signals did not invade a secondary branch (Figures 4D and 4E). In the example shown, the fluorescence response in the left secondary branch (cyan) was lower than that in the parent (yellow) and the right secondary branch (magenta). These differences in peak signal amplitude were statistically significant above 100 Hz (maximum p<0.05, Wallis H-test; Figure 4F), and the differences in area were statistically significant above 120 Hz (maximum p<0.05, Wallis H-test; Figure 4G). Differences in concentration of the calcium indicators or z-plane between branches cannot explain the differences in signal between branches, since at lower frequencies, the response in all branches was the same. This example suggested that, at least in some branching points, the propagation of spikes at high frequencies was not efficient. Similar results were obtained in 5 other cases (see below; Figure 4 - Figure Supplement 1).

**Figure 4.**
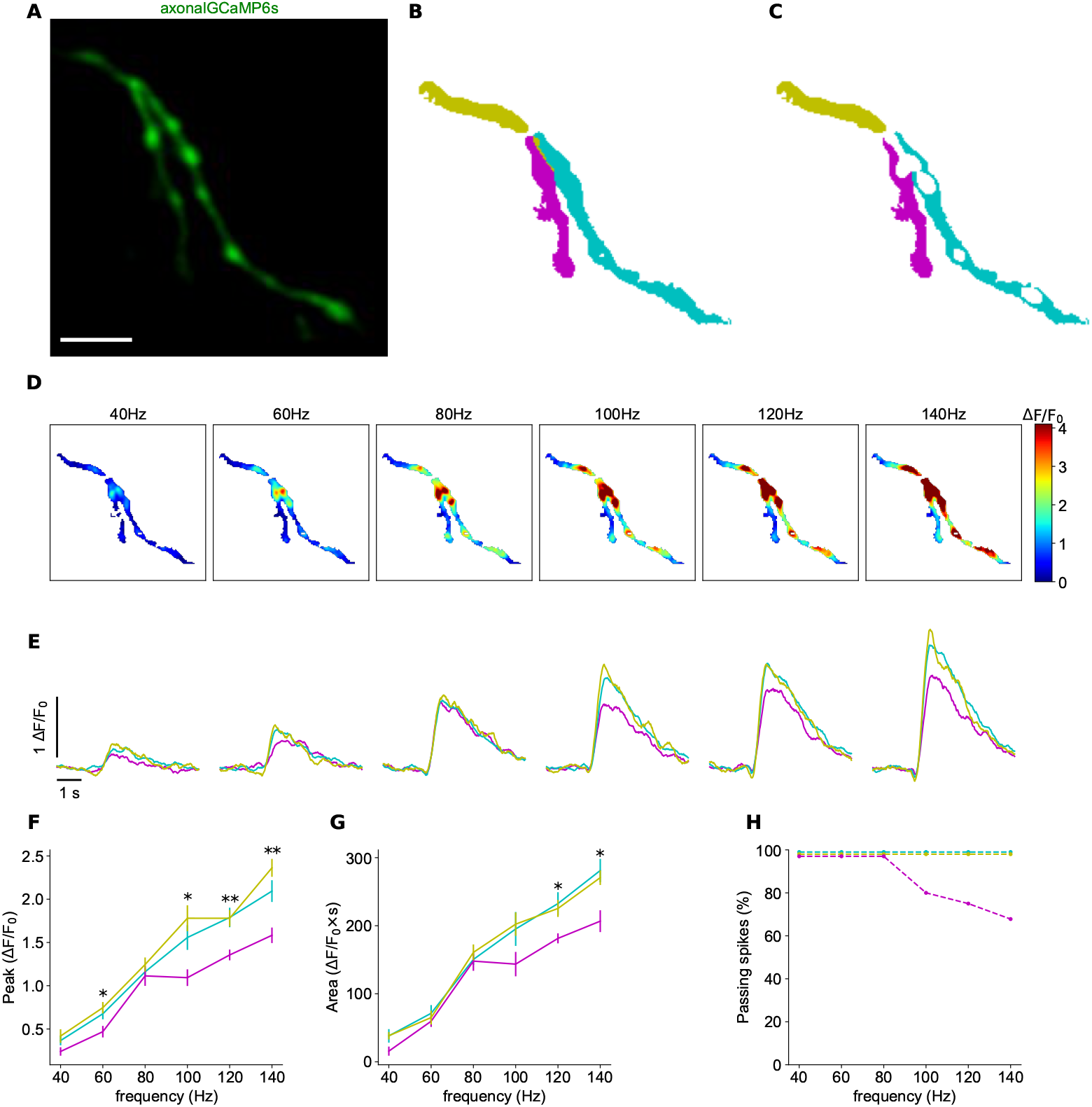
Differential spike propagation in axonal branches. A: Time-averaged image of GCaMP6s activity during electrical stimulation. Scale bar: 5 *μ*m. B: Color map of parent and two secondary branches. C: Color map of parent and two secondary branches after removing the axonal boutons. D: Maps of normalized peak amplitudes at each frequency. E: Normalized GCaMP6s/mRuby3 signal for each branch at different frequency. Average of 7 trials. Colors according to branches in 4C. F: Peak amplitudes of signals in 4E as a function of frequency. G: Area under curve of signals in 4E as a function of frequency. Vertical lines show standard error. Asterisks indicate statistical significance differences between signals; Kruskal-Wallis H-test. H: Percentage of signals that propagate at each branch, as a function of frequency. Figure 4—figure supplement 1. Spike failures at different branchpoints.

### Frequency-dependent filtering of spike propagation

To better understand how spikes propagate or fail, we further analyzed the data from this neuron. The ratio between the action potential train frequency and the peak signal (Figure 4F) and area under the curve (Figure 4G) was linear in the parent branch (yellow) and in the right secondary branch (cyan). This suggested that all spikes of the train propagated through them. Because of the nonlinear relationship between fluorescence and calcium concentration (see Methods), we inferred the number of spikes at the left secondary branch based on the fluorescence intensity in the parent branch.

We estimated the number of spikes that failed to propagate using the area under the curve graphs (Figure 4G), which was more robust to noise than peak of signals (Figure 4F). We found that in one of the secondary branches, the area under the curve was similar at 80 and 100 Hz (148±13.13 Δ*F/F*_0_*s* and 143.58±16.83 Δ*F/F*_0_*s*; flatness in the magenta curve in Figure 4G). We estimated that the same number of spikes – 16 – were propagated in both cases but that only 16 out of 20 spikes (80%) were propagated at 100 Hz. At 120 Hz, 18 out of 24 spikes (75%) propagate into the left daughter (magenta); and at 140 Hz, only 19 out of 28 spikes (67.8%) invaded the left branch (Figure 4H). In conclusion, based on the analysis of one neuron, whereas at lower frequencies all spikes propagate into the two secondary branches, as the frequency of the action potential train increases, the number of spikes that propagated decreased.

### Gradation of spike propagation failures in different neurons

We thus found that spike propagation was effective for most cells, but that there were also cases of propagation failure at high frequency. To assess the extent of propagation failures at all axonal branching points, we designed criteria to classify whether propagation was effective or not and quantified differences in peak and areas of the signals among branches (Figure 5). To achieve this, we calculated the cumulative differences of normalized signals between each pair formed by the parent branch and its two secondary branches (see Figure 5A). These differences increased as the frequency increased and became more pronounced at higher frequencies (see Figures 5C-N). The distribution of signal differences across the 17 examined branchpoints was continuous, although we noted that at 140 Hz, one can distinguish two groups – one with low and one with high peak signal difference values (Figure 5M). The small number of samples (n=17) did not enable statistical tests to test the potential bimodality of the distribution. Our findings also revealed a disparity in signal transmission between the two daughter branches, indicating that, on average, the number of action potentials invading each daughter branch differs, in a gradation of responses.

**Figure 5.**
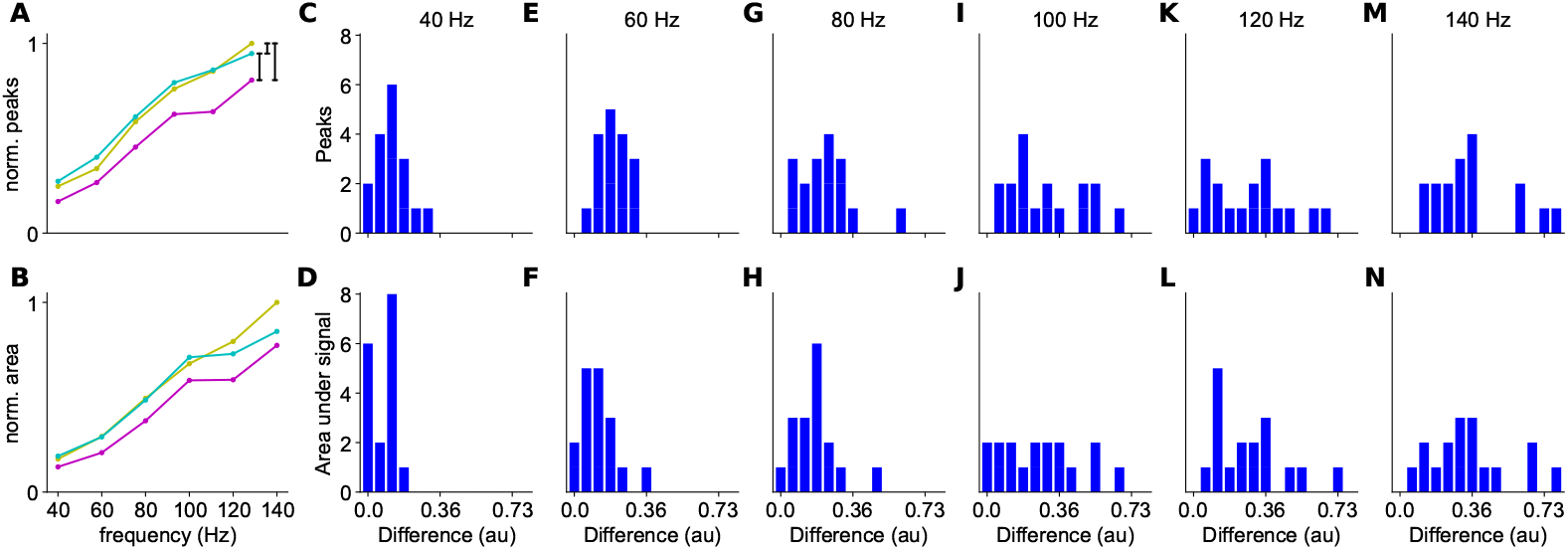
Differences in peak responses and areas between branches. A: Example of the normalized peak fluorescence as a function of stimulus frequency at a branchpoint. Parent branch in yellow, secondary branch with higher signal in cyan, and secondary branch with lower signal in magenta. B: Normalized integrated fluorescence as a function of stimulus frequency. C-N: Differences between normalized signals (black vertical lines in A), for peak (upper row) and area under the signal (bottom row) different spike trains frequencies (n=17).

### Axonal branching point geometry correlates with spike propagation

To further understand the filtering properties of branching points, we compared axonal branch-points that had reliable propagation or filtering properties. For this, we established statistical criteria to distinguish between ‘similar’ and ‘different’ responses in parent and secondary branches. A ‘different’ response was defined if there was a statistically significant difference in peak amplitude or area under the curve in at least two consecutive frequencies (see Figures 4F and 4G). Otherwise, the responses of the branches were classified as ‘similar’ (see Figures 3E and 3F). Using these criteria, we tabulated all the data. In 11 out of 17 samples of axonal bifurcations analyzed we found similar responses in all axonal branches, whereas in the remaining 6 branchpoints, representing data from three different mice, responses were different (Table 1). Next, we measured the diameters of the axonal branches and calculated the geometrical ratio (GR) between the parent and secondary branches, defined as:

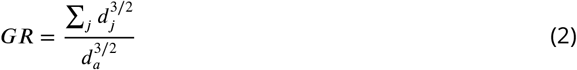

where *d*_*a*_ is the diameter of the parent branch, and *d*_*j*_ are the diameters of the secondary branches. Following cable theory, the GR reflects the electrical impedance at the branching point, correlated with the diameters of the parent and secondary branches. For example, when all branches (parent and secondary) have the same diameter, GR equals 2. Theoretical studies have shown that when GR=1, the propagation of the action potential is effective, whereas higher values of GR may lead to spike delays or failures (***Goldstein and Rall, 1974***; ***Manor et al., 1991***).

We found a significant difference in GR values between the group of branching points with similar responses in all branches compared with the group with different ones (Figure 6A). GR values were lower (1.756 ± 0.328; mean ± std, 1.849; median; p<0.05, t-test; n-neurons=14, n-mice=5;) in the ‘different’ response group than in the ‘similar’ response group (2.304 ± 0.318; mean ± std, 2.149; median). We did not find a difference in angle between the daughter branches between the branching points with and without filtering.

**Figure 6.**
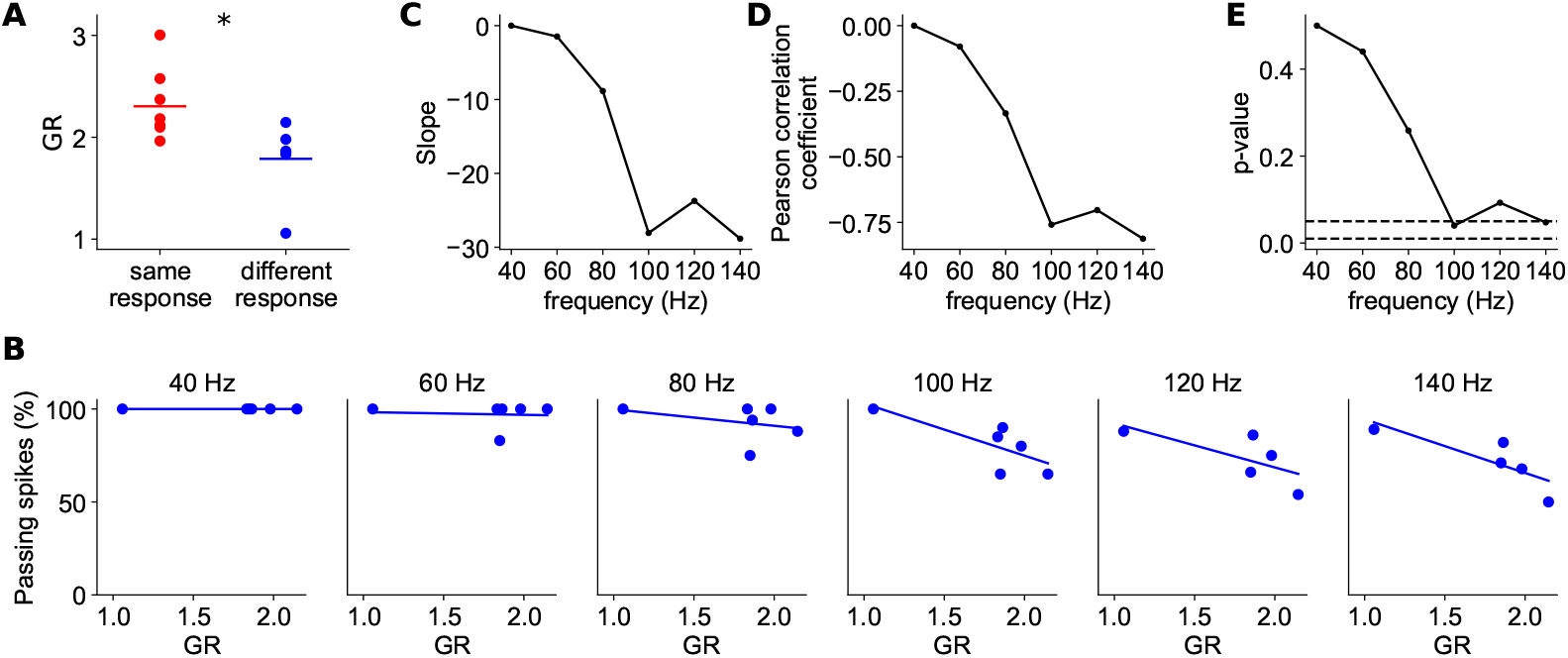
Spike filtering correlates with axonal branchpoint geometrical ratio (GR). A: GR values of ‘similar’ and ‘different’ response branchpoints; t-test two-sided. B: Percentage of propagating spikes as a function of GR for each spike train frequency. Lines represent linear fit to the data. C: Slope of the regression line as a function of action potential frequency. D: Pearson correlation coefficient (R) between the percentage of passing spikes and GR, as a function of action potential frequency. E: p-value of the linear regression fitting as a function of the action potential frequency. Dashed lines indicate p-values of 0.05 and 0.01.

Then, we focused on the subgroup of axonal branchpoints that had ‘different’ responses and tested the correlation between the amount of filtering and GR value. At lower frequencies of 40 and 60 Hz, there was no significant correlation (p>0.4; Wald Test); however, starting at 80 Hz, filtering correlated with GR value (Figures 6B, C, and D). This filtering was higher as the frequency increased, with a statistically significant difference (p<0.05; Wald Test) at 100 and 140 Hz (Figure 6E). We concluded that branchpoints that significantly filtered high-frequency action potential trains had lower geometric ratios and, within this population, the filtering was proportional to GR values.

Interestingly, we also noted that in 6 out of 17 branching points (5/9 cases in the ‘different’ response group and 1/10 cases in the ‘similar’ response group), a bouton was located precisely at the branching point, as can be seen in Figure 2A. This bouton location could potentially impact spike filtering at the branching point due to changes in GR or variations in ion channel densities and types in the bouton membrane.

## Discussion

Here we show the existence of propagation failures of high frequency action potential trains at axonal branchpoints in a significant number of cortical neurons *in vivo*. Axonal computation by axonal branch-specific activity could be important for neuronal information processing, in a similar way to dendritic computation by dendritic branch-specific activity (***Poirazi et al., 2003***; ***Cichon and Gan, 2015***; ***Moore et al., 2022***). Thus, the geometry of the axonal branches may modulate the firing pattern of action potential trains, which is essential for neuronal processing in health and disease. Moreover, axons are also highly sensitive and vulnerable components that are prone to damage, which can further impede spike propagation in cases of traumatic brain injury, stroke, and neurodegenerative diseases such as amyotrophic lateral sclerosis (ALS) or Parkinson’s (***Johnson et al., 2013***; ***Tennant et al., 2017***; ***Gershoni-Emek et al., 2018***; ***Vasu and Kaphzan, 2022***).

### Higher calcium concentration in axonal boutons

Our data demonstrate that axonal boutons have significantly higher calcium accumulations than those in axonal branches. These differences cannot be accounted by differential targeting of the calcium indicator, as while untargeted GCaMP6s expressing axons displayed bright fluorescent varicosities that are likely boutons, axonal targeted GCaMP6s basal fluorescence was relatively homogenous in both axonal shafts and varicosities (***Broussard et al., 2018***). Our results extend the findings of previous *in vitro* reports to the *in vivo* regime. The concentration of calcium in the boutons was reportedly higher than in axonal branches due to other internal stores of calcium (***Emptage et al., 2001***). Bouton responses are variable, as when responses to a single action potential were measured across multiple axonal boutons in the same neuron, there was a more than 10-fold variation in the intensity of calcium transients (***Koester and Sakmann, 2000***). Thus, attempts to use data from a single axonal bouton to infer the firing rate of the presynaptic cell would be prone to error (***Ali and Kwan, 2019***). In our results, we observed a higher calcium concentration at axonal boutons compared to the axonal branches (Figure 2). Therefore, we masked and removed the boutons, enabling us to record the activity from axonal segments without the influence of calcium concentration at the boutons.

### Differential propagation of high-frequency spike trains in different axons

Importantly, and different from previous reports *in vitro*, we found two types of branchpoint propagation. In the first group, all spikes reliably invaded all axonal branches, while in the second group, there was a significant failure of spike propagation at high frequencies. Interestingly, GR values in the ‘different’ response group, where some spikes failed to propagate, were lower than GR values in the ‘similar’ response group (Figure 6A). But, by independently analyzing the ‘different’ response group, we found a correlation between the percentage of spike filtering and GR (Figure 6B), as expected by cable theory. This correlation is inconsistent with the group of branching points with higher GR values, that present similar activity in all branches, without spike failures. Thus, besides cable properties, additional mechanisms must be considered to elucidate the different phenomenology between these two groups. These differences may be attributed to variations in ion channel types and their respective densities. Also, these subgroups belong to different neuronal subtypes, each characterized by distinct membrane properties. In the branching points of the ‘similar’ response group, it is possible that higher values of GR are required to induce spike failures.

### Comparison with previous studies

Previous studies conducted in brain slices did not report spike failures in axonal bifurcations with relative lower frequency trains. Most of the spike filtering we report was detected at higher frequencies of above 100 Hz. While typical frequencies of spike trains in neocortical pyramidal neurons at normal conditions are below 50 Hz (***Quiroga et al., 2009***; ***Apostolides et al., 2016***), higher frequencies of up to 150 Hz have been observed *in vivo* under physiological conditions (***De Kock et al., 2007***; ***Larkum et al., 2007***). In addition, previous studies *in vitro* examined branchpoints located in close proximity to the soma, typically within the first 300 *μ*m (***Foust et al., 2010***; ***Popovic et al., 2011***). This is often where the primary collaterals of the axon originate (***Debanne, 2004***). But it is likely that spike failures might tend to be more prevalent in more distal axonal arbors, like the one we studied. In our experiments, while precise measurements of the distance between the branching point and the cell body were not feasible due to anatomical constraints, we estimate that the axonal branching points are located at least 1 mm away from the electrode and the viral expression site. Furthermore, it is worth noting that axonal regions near the soma tend to have a higher degree of myelination (***Tomassy et al., 2014***), which may prevent spike failures (***Hamada et al., 2017***). Our experimental setup does not enable us to see the presence or absence of myelin sheath around the axons. Previous evidence show that most of the axons are unmyelinated, mainly in distal regions. In the mouse somatosensory cortex, the amount of myelinated axons in layer 2/3 is around only 37%, lower than in layer 4 (56.7%) and layers 5/6 (63%) (***Tomassy et al., 2014***). In a reconstruction of human temporal cortex, 40.6% of the volume was unmyelinated axons and 7.6% was myelinated axons (***Shapson-Coe et al., 2021***), meaning that only ∼16% of the axons are myelinated. Additionally, previous calcium measurements were contaminated by axonal boutons (***Cox et al., 2000***), which typically exhibit higher and more variable calcium concentrations.

### Potential mechanisms and functional consequences

Our results are consistent with the possibility that spike filtering is partly due to the electrical cable structure of the axon, as determined by its geometry, as predicted by Rall (Figure 6). In addition, in some branchpoints we found axonal boutons at the point of bifurcation (for example see Figure 2A). These boutons can affect spike filtering, due to their morphology as well as the ion channel types and densities on the bouton membrane. In addition to geometrical factors of the axonal bifurcations, other mechanisms, such inhibitory axons that form axo-axonic synapses onto the axonal tree, may cause action potential failures and spike filtering. For example, cholinergic Kenyon cells in Drosophila have numerous axo-axonic connections that suppress signals in neighboring cells (***Manoim et al., 2022***). GABAergic neurons in the spinal cord of rodents and humans also form axo-axonic contacts with sensory axons, facilitating spike propagation by preventing spike failures at axon branchpoints (***Hari et al., 2022***). Excitatory synaptic inputs into dopaminergic axons were also found in the mouse striatum, suggested as a physiological mechanism to regulate dopamine signaling (***Liu et al., 2022***).

In addition, different anatomical characteristics in some axons are consistence with the specific spike propagation properties within the axonal tree. Along the axon, afferent synapses are closer to the soma than the efferent synapses (***Tomassy et al., 2014***), and the presynaptic axonal boutons target inhibitory neurons in more proximal locations along the axonal tree than the boutons that target excitatory neurons (***Schmidt et al., 2017***). Additionally, boutons in individual axons that simultaneously innervate motor and sensory areas of the cerebral cortex present significant area-specific differences in size (***Rodriguez-Moreno et al., 2020***). Neurons with axons emerging from dendrites rather than from soma were suggested to enable information gating (***Hodapp et al., 2022***). Finally, a neuron with two separate axons emerging from the soma was found in the human temporal cortex (***Shapson-Coe et al., 2021***).

### Dynamic morphological changes of axons

As we have demonstrated, the geometry of the axon, especially the diameters at branching points, has a significant influence on spike filtering. Previous studies showed activity-dependent plasticity of axonal diameters and bouton sizes that affect action potential conduction velocity (***Chéreau et al., 2017***; ***Griswold et al., 2023***). Here we suggest that these dynamical changes in axonal dimensions may affect not only the velocity of action potential propagation but also filtering properties. This dynamical spike filtering ability could have implementations in learning and memory.

Our results are but a ‘proof of concept’ that some spikes of high-frequency action potential trains generated by an extracellular electrode may fail at axonal branching points. More natural scenarios of high-frequency spike trains can be examined in future work by specific visual, auditory, or air puff stimuli. Higher temporal resolution, at the level of a single action potential, should also be obtained in future studies, perhaps by using newly engineered voltage sensors designed for recording axonal activity.

## Acknowledgments

We would like to thank Yuriy Shymkiv and lab members for their support and helpful suggestions; This work was supported by NEI (R01EY011787), NIMH (R01MH115900), NINDS (RM1NS132981) to R.Y.

## Author Contributions

N.O., V.C., and R.Y. contributed to project conception, project design, and manuscript writing. V.C. performed the experiments and N.O. analyzed the results. R.Y. directed the project and secured resources and funding.

## Competing Interests statement

The authors declare no competing interests.

---

**Figure 3—figure supplement 1.**
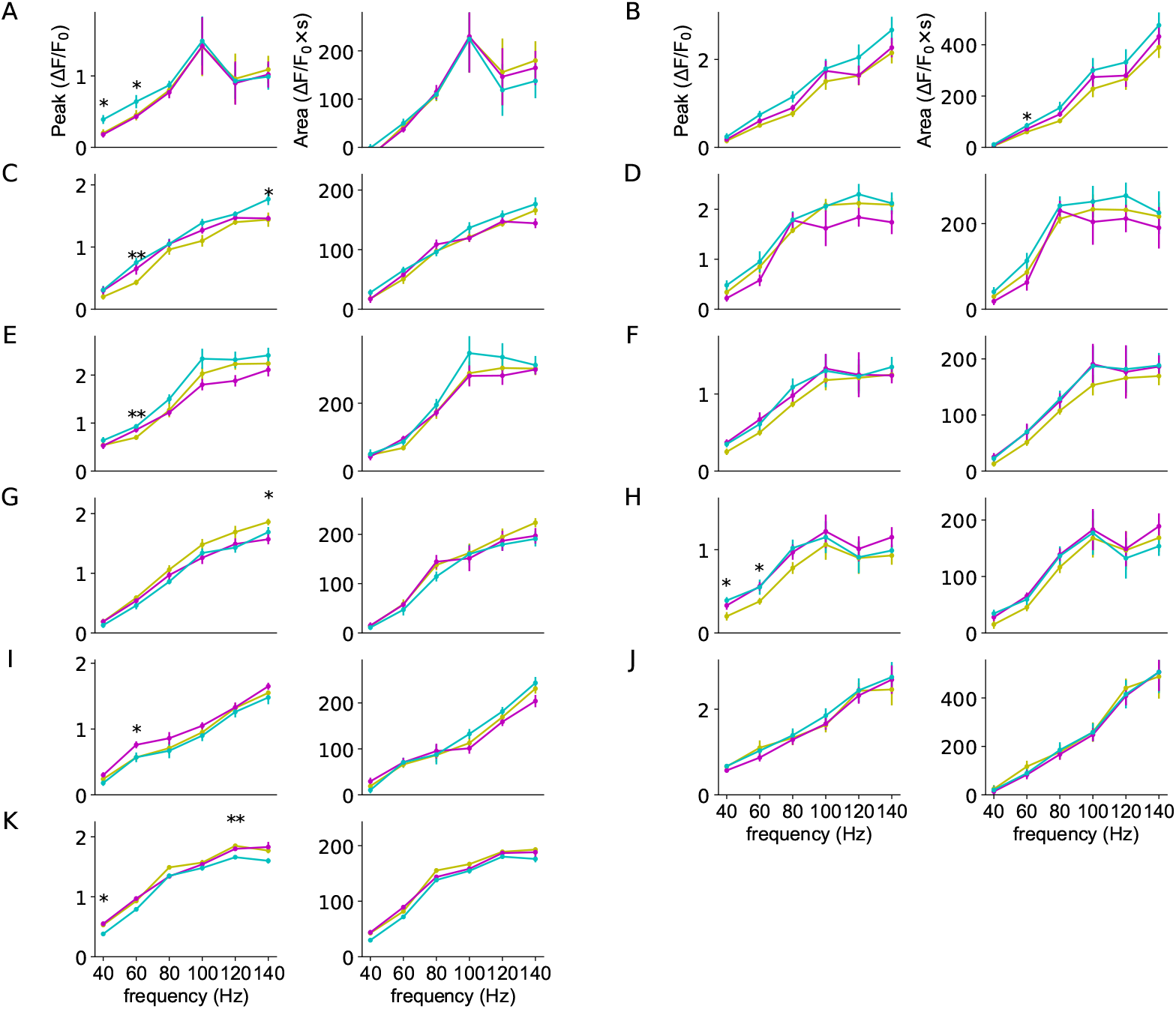
Spike propagation at different branchpoints. A-J: Examples of reliable propagation in both peak (left columns) and integrated fluorescence (right columns). Statistical definition of propagation or failure explained in text. Parent branch in yellow, secondary branches in cyan and magenta. Vertical lines show standard error. Asterisks indicate statistical significance differences between the three branches; Kruskal-Wallis H-test.

**Figure 4—figure supplement 1.**
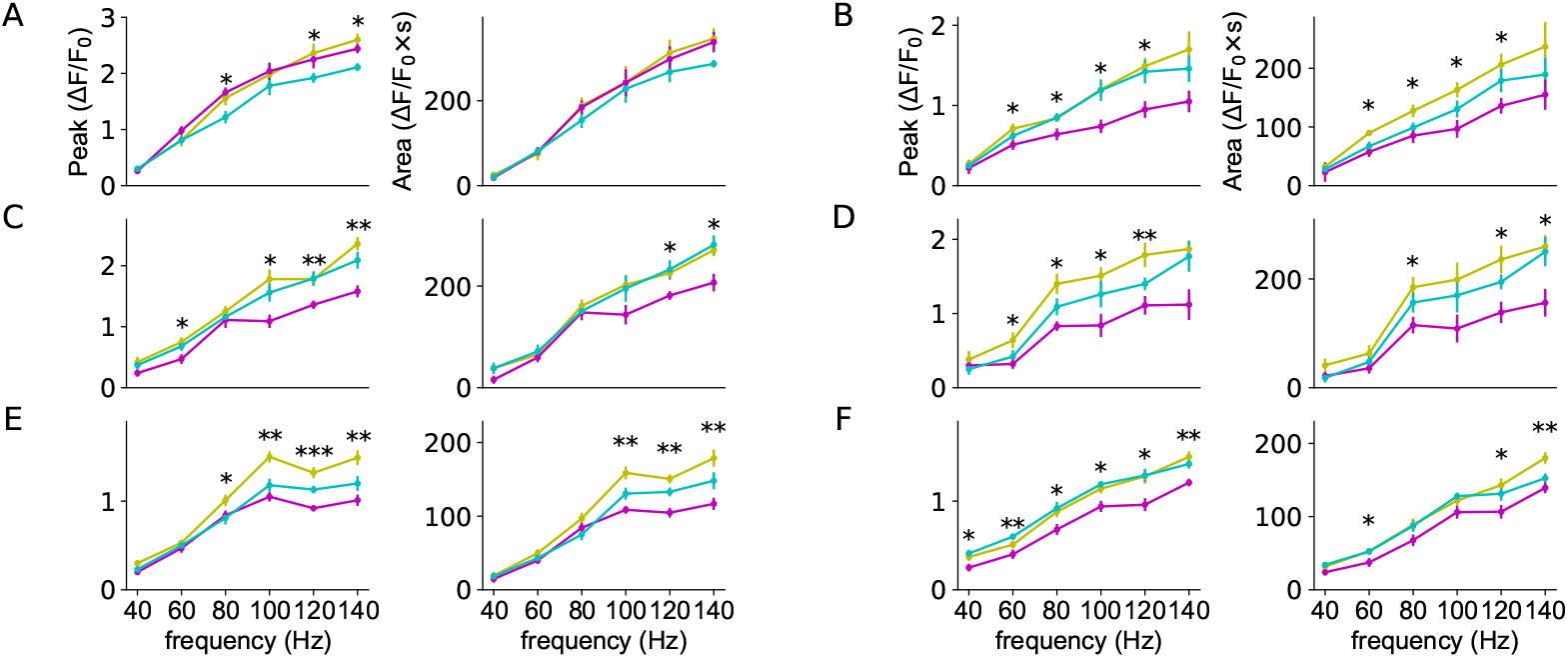
Spike failures at different branchpoints. A-F: Examples of High-frequency propagation failures in both peak (left columns) and integrated fluorescence (right columns). Statistical definition of propagation or failure explained in text. Parent branch in yellow, secondary branches in cyan and magenta. Vertical lines show standard error. Asterisks indicate statistical significance differences between the three branches; Kruskal-Wallis H-test.

## Notes

### Competing Interest Statement

The authors have declared no competing interest.

